# Orally administered antigen can reduce or exacerbate pathology in an animal model of inflammatory arthritis dependent upon the timing of administration

**DOI:** 10.1101/2022.01.07.475399

**Authors:** Gavin R Meehan, Iain B McInnes, James M Brewer, Paul Garside

**Author notes:** **Corresponding Author:** Paul Garside, Institute of Infection, Immunity & Inflammation, College of Medical, Veterinary & Life Sciences, Sir Graeme Davies Building, University of Glasgow, 120 University Place, Glasgow G12 8TA, E.

## Abstract

Currently, treatments for rheumatoid arthritis (RA) are focussed on treatment of disease symptoms rather than addressing the cause of disease, which could lead to remission and cure. Central to disease development is the induction of autoimmunity through a breach of self-tolerance. There is considerable research in RA focussed on antigens and approaches to re-establish antigen specific tolerance. A crucial step in this research is to employ appropriate animal models to test prospective antigen specific immunotherapies, preferably in the context of joint inflammation.

In this short communication, we use our previously developed model of antigen specific inflammatory arthritis in which OVA-specific TcR tg T cells drive breach of tolerance to endogenous antigens to determine the impact that the timing of therapy administration has upon disease progression. Using antigen feeding to induce tolerance we demonstrate that administration prior to articular challenge results in a reduced disease score as evidenced by pathology and serum antibody responses. By contrast, feeding antigen after articular challenge had the opposite effect and resulted in the exacerbation of pathology.

Although preliminary, these data suggest that the timing of antigen administration may be key to the success of tolerogenic immunotherapies. This has important implications for the timing of potential tolerogenic therapies in patients.

## Introduction

Rheumatoid arthritis (RA) is a chronic inflammatory condition in which a series of genetic and environmental factors trigger a breach of immunological self-tolerance. This results in the development of autoimmunity and ultimately culminates in the destruction of the bone and cartilage of the joints. Current therapies for the treatment of RA focus on decreasing inflammation and preventing disease progression but do not treat the underlying cause of pathology. Consequently, there has been a drive for recent arthritis research to focus on the development of antigen-specific immunotherapies that would restore immunological homeostasis and allow for drug-free remission [1].

Tolerogenic therapies can take many forms including tolerogenic dendritic cells (tolDCs), regulatory T cell (Treg) induction, tolerogenic liposomes and antigen feeding [2], Various pre-clinical models have examined these therapies and a number are undergoing clinical trials for the treatments of other autoimmune diseases [3–5]. However, questions remain regarding the form these therapies should take and how they should be administered. In particular, the timing of administration is key to ensuring that the therapies are not only successful but do not exacerbate disease. Thus, will tolerogenic therapies require administration in at risk patients prior to overt pathology or will they be effective in the latter scenario?

To examine the impact of timing on antigen-specific immunotherapy administration we employed an ovalbumin (OVA) induced model of antigen specific inflammatory arthritis, in which OVA-specific TcR tg T cells drive breach of tolerance to endogenous antigens. Using this model, we fed mice OVA protein at various stages of disease to determine the impact of antigen feeding on pathology and whether tolerance could be induced.

## Materials and Methods

### Acknowledgements

We acknowledge the assistance of the Histology Research Service at the University of Glasgow. We also acknowledge Dr Robert Benson and Veronica Patton at the University of Glasgow for their assistance in scoring histology sections.

### Funding

This project has received funding from the Innovative Medicines Initiative 2 Joint Undertaking under grant agreement No 777357. This Joint Undertaking receives support from the European Union’s Horizon 2020 research and innovation programme and EFPIA. www.imi.europa.eu

### Disclaimer

This communication reflects the authors views and neither IMI nor the European Union, EFPIA, or any Associated Partners are responsible for any use that may be made of the information contained within.

### Animals

C57BL/6J mice were purchased from Envigo (Wyton, UK). CD45.1*OTII mice were produced in house (Central Research Facility, University of Glasgow, UK). Animals were maintained on a 12-hour light/dark cycle and provided with food and water ad libitum. All procedures were performed under a UK Home Office licence in accordance with the Animals (Scientific Procedures) Act 1986.

### Induction of OVA Breach of Tolerance Arthritis Model

The OVA breach of tolerance arthritis model was used as described previously [6,7]. Briefly, CD4 T cells were isolated from the lymph nodes and spleens from 6-12-week-old OTII mice. Th1 cell differentiation was induced by culturing CD4 T cells with antigen presenting cells treated with 50μg/ml mitomycin C (Merck, Darmstadt, Germany) in the presence of 1μg/ml OVA^323-339^ (Peprotech, Rockyhill, NJ, USA), 10ng/ml IL-12 (R&D systems, Minneapolis, MN, USA) and 2μg/ml anti-IL4 (Biolegend, San Diego, CA, USA) for 3 days. The purity of the OTII cells was confirmed by flow cytometry and 3,000,000 cells were injected intravenously into recipient C57BL/6J mice. The following day mice were injected subcutaneously with 100μl of 100μg OVA emulsified in Freund’s complete adjuvant (CFA) (Sigma Aldrich, St Louis, MO, USA). 21 days later the mice were challenged with a periarticular injection of 50μl PBS containing 100μg heat aggregated OVA (HAO) into a hindlimb. Rechallenges, when performed, consisted of an articular injection of 50μl Freund’s incomplete adjuvant (IFA) (Sigma Aldrich, St Louis, MO, USA) containing 100μg OVA given 63 days after the first HAO challenge. Mice were weighed and monitored daily for signs of arthritis. Each footpad was measured using digital callipers and given a disease score based upon erythema, swelling and loss of function as described previously [6].

### Antigen Feeding

Ovalbumin protein (Sigma Aldrich, St Louis, MO, USA) was prepared in sterile water at 40mg/ml and gently agitated at 4°C overnight. The dissolved ovalbumin was filtered through a 0.22μM membrane and added to sterile water bottles in the treatment groups cages for 10 days. The water bottles were changed daily. Control mice received sterile tap water. The timing of antigen feeding is indicated in each experiment.

### Histology

Histology was performed as described previously [8]. Briefly, hind limbs were collected and stored in 10% neutral buffered formalin. The tissue was then decalcified in 5% formic acid and processed for wax embedding. Tissue sections (8μm) were cut along the sagittal plane and stained with haematoxylin and eosin or toluidine blue. Images were taken using an EVOS Cell Imaging System (Thermofisher, Waltham, MA, USA). Scoring was performed by a blinded observer based on a scale of 0-3 for cellular infiltration, synovial hyperplasia and cartilage/bone erosion as described previously [6]. In addition, mice were given a score of 0 or 1 based upon the presence of ulceration. This provided each mouse with a total score out of 10.

### Serum Antibody ELISA

The levels of serum anti-OVA or anti-collagen type ii (CIi) IgG1 and IgG2C were measured using enzyme linked immunosorbent assays (ELISA) as described previously [6]. Briefly, ELISA plates (Corning Inc, Corning, NY, USA) were coated with 20μg/ml OVA protein (Sigma Aldrich, St Louis, MO, USA) or 4μg/ml CII (Sigma Aldrich, St Louis, MO, USA) in sodium bicarbonate buffer (Sigma Aldrich, St Louis, MO, USA) overnight at 4°C. Plates were washed in PBS-T and blocked in animal free block (Vector Laboratories, Burlingame, CA, USA) for 1 hour at 4°C. Serum samples were prepared at 1:50 and serially diluted across the ELISA plate. The samples were incubated overnight at 4°C. The plates were then washed in PBS-T. Biotin anti-mouse IgG1 or IgG2C (Jackson Laboratory, Bar Harbor, ME, USA) were prepared in PBS at 1:5000 and 1:2000 dilutions respectively and added for 1 hour at 4°C. The plates were washed again in PBS-T and incubated with ExtrAvidin peroxidase (Sigma Aldrich, St Louis, MO, USA) at a 1:10000 dilution for 1 hour at 4°C. The plates were washed in PBS-T and developed using SIGMAFAST OPD tablets ((Sigma Aldrich, St Louis, MO, USA) in the dark at room temperature for 20 minutes. The plates were stopped with the addition of 50μl of 10% sulfuric acid and then read at 492nm using a Tecan ELISA plate reader (Tecan Group, Männedorf Switzerland).

### Statistical Analysis

All graphs and statistical analyses were produced using GraphPad Prism 7 (GraphPad Software Inc., San Diego, CA, USA). P values < 0.05 were deemed to be significant.

## Results and Discussion

To determine whether feeding antigen could induce tolerance we used the OVA breach of tolerance model of inflammatory arthritis. Using this model, we fed OVA to mice either pre or post immunisation with OVA/CFA or post articular challenge with HAO (Fig 1a). Feeding OVA before OVA/CFA immunisation resulted in a significant reduction in footpad swelling 24 hours post HAO challenge (Two-Way ANOVA, **<0.01) (Fig 1b). This was consistent with historic antigen feeding studies where animals fed soluble type II collagen before the induction of collagen induced arthritis (CIA) experienced less severe disease with a delayed onset [9,10].

**Figure 1.**
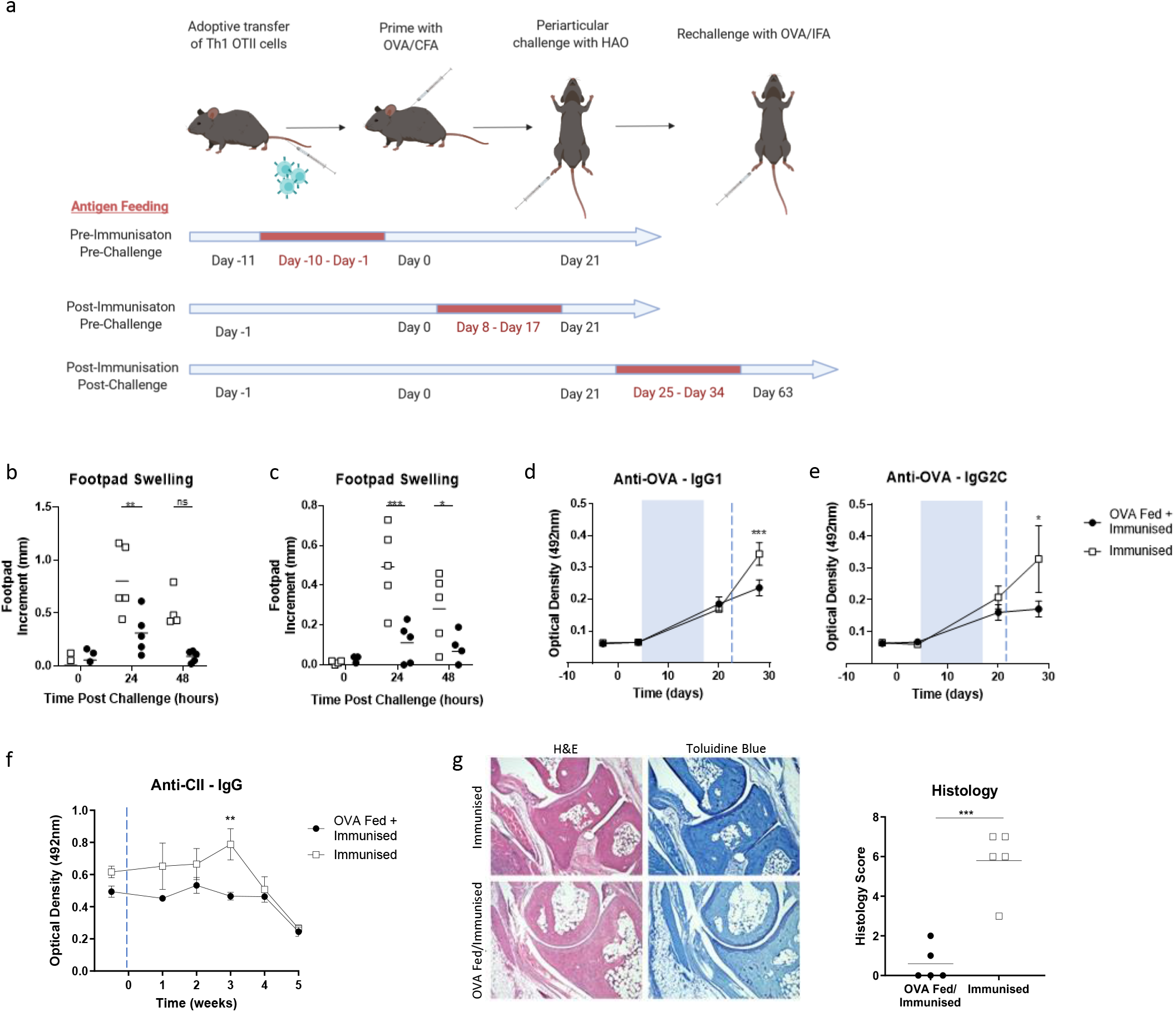
Antigen feeding pre-challenge induces tolerance. (a) The OVA model of inflammatory arthritis was used to study tolerance. CD4 T cells that specifically recognise OVA^323-339^ in the context of MHCII were adoptively transferred into C57BL/6 mice. The mice were immunised with OVA/CFA and were then given an articular challenge of heat aggregated OVA (HAO) in PBS. Rechallenges, when performed, consisted of an articular challenge with OVA/IFA. Antigen feeding was performed by supplementing drinking water with ovalbumin for a ten-day period (as indicated in red) in separate experiments. Control mice were not fed antigen. Footpad measurements were taken of mice that had been fed ovalbumin pre (b) and post (c) - immunisation with OVA/CFA. Measurements were taken 0, 24 and 48 hours post HAO challenge. n=5 from one independent experiment. Statistical analysis was performed using a Two-Way ANOVA, ns = no significance, *<0.05, **<0.01, ***<0.001. ELISAs were performed on the serum of mice that had been fed ovalbumin postimmunisation. These examined anti-OVA IgG1 (d), anti-OVA IgG2C (e) and anti-CII IgG (f) antibodies. n=5 from one independent experiment. Statistical analysis was performed using a Two-Way ANOVA, *<0.05, ***<0.001. Blue shaded boxes signify period of antigen feeding. Blue dashed line signifies HAO challenge. (g) Histology was performed on the joints of mice fed ovalbumin post-immunisation. n=5 from one independent experiment. Disease scoring was performed blinded. Statistical analysis was performed using a n unpaired t test, ***<0.001. Figure 1a created with BioRender.com.

A similar effect was observed following OVA feeding after immunisation with OVA/CFA (Fig 1c). Footpad swelling was significantly reduced both 24 hours (Two-Way ANOVA, *<0.05, ***<0.001) and 48 hours (Two-Way ANOVA, *<0.05) post HAO challenge. This was accompanied by a significant reduction in anti-OVA IgG1 (Two-Way ANOVA, ***<0.001) (Fig 1d) and IgG2C (Two-Way ANOVA, *<0.05) (Fig 1e) antibodies in the OVA fed group. Similarly, anti-collagen II (CII) antibodies that were tracked for several weeks post HAO challenge were consistently lower in the OVA fed group although there was only a significant difference at week 5 (Two-Way ANOVA, **<0.01) (Fig 1f). The histology score, based upon cellular infiltration, synovial hyperplasia, and cartilage/bone erosion, was also significantly reduced in this group (Unpaired t test, ***<0.001) (Fig 1g). These results were similar to those found with models that combine CIA and a second protein such as OVA to induce arthritis in CIA resistant mice. As with our model, feeding these mice OVA before or after disease induction significantly reduced footpad swelling and articular inflammation [11].

Having demonstrated that OVA feeding prior to breach of tolerance was able to reduce disease severity we next fed antigen to mice following HAO challenge. To best measure the effects of antigen feeding at this time point we re-challenged the mice with OVA/IFA. Although there was no significant difference in footpad swelling 24 hours post re-challenge (Fig 2a), subsequent time points indicated a significant increase in footpad swelling in the OVA fed/re-challenged group that became progressively greater over time (Two-Way ANOVA, **<0.01, ****<0.0001). Due to the extent of this swelling in the OVA fed/re-challenged mice the experiment was terminated. The severity of the pathology is apparent in Fig 2b where substantial swelling, erythema and ulceration is present in the OVA fed/re-challenged group compared to the re-challenged group. Examination of the serum found no significant increase in anti-OVA IgG1 antibodies over time (Fig 2c) but did show a significant increase in anti-OVA IgG2C antibodies at day 45 following OVA feeding but pre-rechallenge (Two-Way ANOVA, ***<0.001) (Fig 2d). There were no significant differences in anti-CII antibodies at any time point between the two groups (Two Way ANOVA) (Fig 2E). Interestingly, there was no difference between the treatment groups at the final time point with any of the antibodies we examined suggesting that the observed effect may not be antibody-mediated. It is possible that another autoantibody that we have not measured may be driving inflammation, but the low serum anti-OVA and anti-CII antibodies suggests that the T cells themselves may be directly mediating inflammation by driving recruitment of innate immune cells into the joint as indicated in the histology.

**Figure 2.**
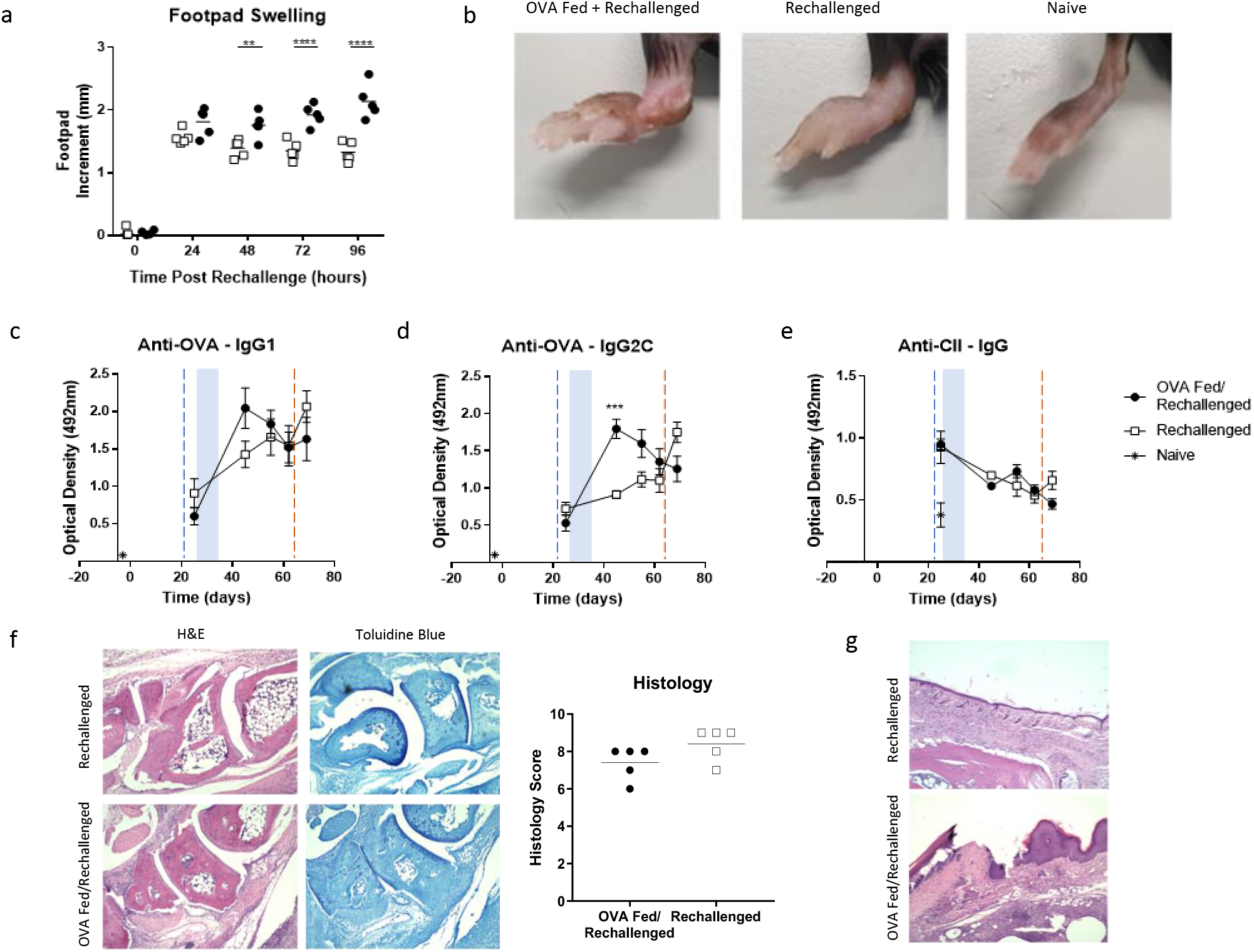
Antigen feeding post-challenge exacerbates disease. (a) Following induction of inflammatory arthritis, mice were fed ovalbumin following a challenge with heat aggregated ovalbumin (HAO). Control mice were not fed antigen. The mice were then rechallenged with OVA/IFA and footpad measurements were taken 0, 24, 48, 72 and 96 hours post rechallenge (a). n=5 from one independent experiment. Statistical analysis was performed using a Two-Way ANOVA, ns = no significance, **<0.01, ***<0.0001. (b) Representative photographs of the challenged footpads indicate differences in the disease states of the mice. ELISAs were performed on the serum of mice that had been fed ovalbumin post-immunisation. These examined anti-OVA IgG1 (c), anti-OVA IgG2C (d) and anti-CII IgG (e) antibodies. n=5 from one independent experiment. Statistical analysis was performed using a Two-Way ANOVA, ***<0.001. Blue shaded boxes signify period of antigen feeding. Blue and red dashed lines signify HAO challenge and OVA rechallenge respectively. (f) Histology was performed on the joints of mice fed ovalbumin postimmunisation. n=5 from one independent experiment. Disease scoring was performed blinded. Statistical analysis was performed using a n unpaired t test. (g) Histological images indicate the presence of ulceration in the footpads of the OVA fed/rechallenged mice.

Histological examination of the joints showed substantial bone erosion, hyperplasia, and cellular infiltration in both the re-challenged and OVA fed/re-challenged mice. Although there was no significant difference in the histology scores (Unpaired t test) (Fig 2F), we hypothesise that the severity of the inflammation in both treatment groups may have obscured our ability to distinguish subtle differences between them. One notable difference was the presence of substantial ulceration in the footpads of the OVA fed/re-challenged mice (Fig 2G). The presence of ulceration suggests a more severe inflammatory response in these mice. This could be due to the antigen feeding expanding effector T cells or the absence or loss of Treg suppression, but a detailed immunological analysis would need to be performed to assess this further.

Taken together, these data suggest that earlier interventions with tolerogenic therapies may be key to their success. Many previous animal studies in both immunisation [12,13] and disease models [14,15] have demonstrated that it is relatively easy to induce tolerance prophylactically whereas it is much more challenging to tolerise an already primed immune response. Critically, studies in a murine model of autoimmune diabetes found an exacerbation of pathology when attempting to tolerise a primed immune response [16].

Although it is unclear if these results translate into human disease, they suggest that tolerogenic therapies would be best targeted at individuals at risk of developing or in the very early stages of RA. In contrast, attempting to tolerise individuals in the clinical phase of RA may result in an exacerbation of symptoms and a poorer outcome. In addition, we have recently demonstrated in our animal model that the repertoire of antigens to which tolerance is breached becomes wider at later time points following initial breach of tolerance providing another reason to target therapy early [17]. Further work should be performed to determine whether antigen feeding can induce tolerance in drug controlled clinical phase arthritis and whether other tolerogenic therapies are also dependent upon the timing of administration.

